# A class of identifiable phylogenetic birth-death models

**DOI:** 10.1101/2021.10.04.463015

**Authors:** Brandon Legried, Jonathan Terhorst

## Abstract

In a striking result, Louca and Pennell (2020) recently proved that a large class of phylogenetic birth-death models are statistically unidentifiable from lineage-through-time (LTT) data: any pair of sufficiently smooth birth and death rate functions is “congruent” to an infinite collection of other rate functions, all of which have the same likelihood for any LTT vector of any dimension. As Louca and Pennell argue, this fact has distressing implications for the thousands of studies that have utilized birth-death models to study evolution.

In this paper, we qualify their finding by proving that an alternative and widely used class of birth-death models is indeed identifiable. Specifically, we show that piecewise constant birth-death models can, in principle, be consistently estimated and distinguished from one another, given a sufficiently large extant time tree and some knowledge of the present-day population. Subject to mild regularity conditions, we further show that any unidentifiable birth-death model class can be arbitrarily closely approximated by a class of identifiable models. The sampling requirements needed for our results to hold are explicit, and are expected to be satisfied in many contexts such as the phylodynamic analysis of a global pandemic.

## 1 Introduction

The birth-death process (Feller, 1939; Kendall, 1948) is a classic model of population growth. Recently, it has also been used to study the processes of speciation and extinction (Nee et al., 1994; Morlon et al., 2010, 2011; Morlon, 2014), and also the evolution of pathogens (Stadler et al., 2013). Data-driven inquiry in these fields is inherently challenging because the majority of species and pathogens that have ever lived have left no record of their existence. Thus, we can only make inferences about evolution on the basis of a biased sample of the species or lineages that happened to survive to the present day (Quental and Marshall, 2010; Morlon, 2014). Interest in the birth-death process arises in part from the fact that it provides a principled way of correcting this bias (Höhna et al., 2011; Cusimano et al., 2012).

Realizations of the birth-death process can be viewed from a phylogenetic perspective as rooted trees, where each leaf node represents a species that survived until the present, internal nodes are unobserved species, and edges represent lines of descent. The shape of the tree is governed by two non-negative functions which describe, at any given time *t* before the present, the per-capita rates of birth and death. As noted above, a distinguishing feature of this model is that lineages that died out before the present are not reflected in the resulting tree. Given birth and death rates, as well as a third parameter known as the sampling rate, we refer to the resulting distribution over random trees as a phylogenetic birth-death (henceforth, BD) model. (A precise definition is given in the next section.) The BD model implies a distribution over observed evolutionary data, and given such data, we can use statistical estimation to make inferences about the model parameters.

BD models have been utilized in thousands of published studies (McPeek and Brown, 2007; Condamine et al., 2019; Henao Diaz et al., 2019), despite possessing known and somewhat troubling limitations. Stadler (2009) showed there exist different birth-death models that have the same likelihood in terms of observable data. In statistical terms, this implies that the BD model is *unidentifiable* without further assumptions. The models considered by Stadler are highly parsimonious, consisting of constant birth and death rates that do not change over time. The problem is made even more challenging if the rates are time-varying (Gavryushkina et al., 2014).

Very recently, Louca and Pennell (2020, cited hereafter as LP) proved that the situation is much worse than was previously realized: for *any* reasonably smooth birth and death rate functions, there are *infinitely* many other such functions that result in the same likelihood over phylogenetic trees. Although each of these functions represents a qualitatively different evolutionary scenario, LP‘s result shows that it is impossible to tell which of them produced a given data set, even if the data were infinite. In light of the huge number of times that this model has appeared in the literature, this finding is highly worrisome. There they constructed a pulled (birth) rate function that is identifiable from LTT data.

Consistent estimation is impossible in an unidentifiable statistical model, so when faced with one, there are two ways forward: a) use a different model, or b) impose additional regularity conditions on the parameter space to restore identifiability. For the BD model, option a) may be warranted in some settings, but such a debate is beyond our scope. In this paper, we focus on option b). Our main result is to prove that there exists a class of BD models which are identifiable based on LTT data from an extant time tree. By identifiable, we mean that, within the space of rate functions we consider, each distinct BD model corresponds to one and only one likelihood function, and conversely. In fact, this space consists simply of piecewise constant functions, which are already widely used to fit BD models in practice.

Our results show that this class is identifiable once there are enough leaves in the extant tree, and we derive explicit lower bounds on the requisite number of samples. These bounds depend on a measure of parsimony of the underlying model class: they require that identifiable classes of birth-death rate functions do not oscillate unnaturally, in a sense which is made precise below. The same phenomenon has previously been noticed in population genetics (Myers et al., 2008; Bhaskar and Song, 2014), and our proofs are based in part on these earlier works.

## 2 Preliminaries

In this section, we define the BD model and introduce some key definitions.

Throughout the paper, *n* is used to denote sample size. We assume *n* ≥ 2 and suppress explicit dependence on it when there is no risk of confusion. Given *n* sampled taxa, an *extant timetree* is a bifurcating tree that traces out the ancestry of the sample. Therefore, the extant timetree has *n* − 1 internal nodes which denote the times at which various taxa diverged from common ancestors. These are denoted 0 ≤ *τ*_*n*−1_ *<*· · · *< τ*_1_, where time runs backwards from the present. As in LP, we assume that all *n* samples are collected at time *t* = 0. There is also a root node referred to as the *origin* which occurs at height *τ*_*o*_ *<* ∞, when the process is assumed to have started. Note that the height of the origin node is not resolvable from character data evolving along the tree since it is ancestral to the entire sample. Its value is therefore conditioned on using prior information. An example of an extant time tree with sample size three is shown in Figure 1.

**Figure 1:**
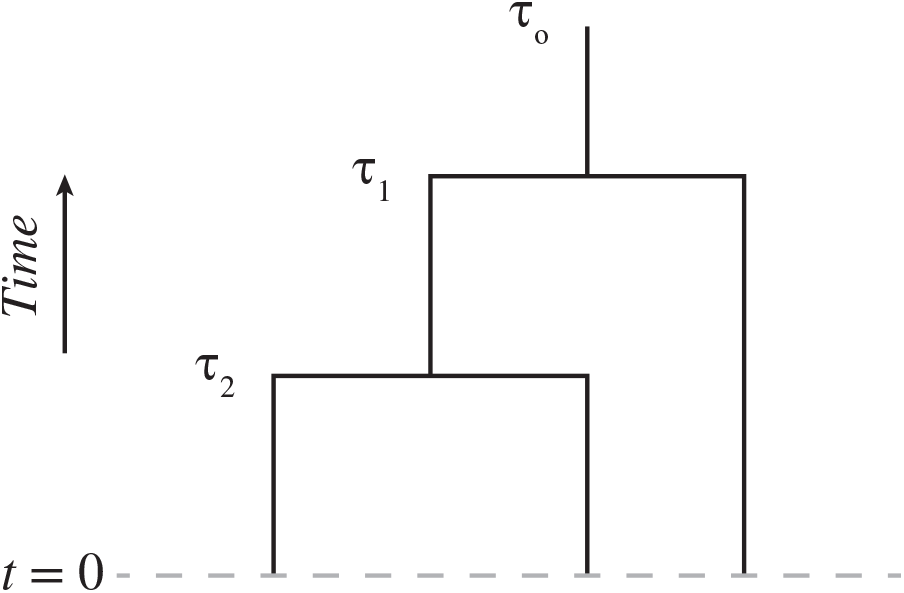
An extant timetree on *n* + 1 = 3 leaves.

Extant timetrees are assumed to be stochastically generated by a BD process (Morlon et al., 2010). This process has three parameters: two positive rate functions *λ*: ℝ _≥0_ → ℝ_*>*0_ and *μ*: ℝ _≥0_ → ℝ_*>*0_, and an initial sampling fraction *ρ* ∈ (0, 1]. Here *λ* and *μ* represent the instantaneous rate per capita at which lineages are born and die going forwards in time, and *ρ* is the fraction of extant lineages that are sampled at the present time *t* = 0. In the sequel, we refer to different BD models by their corresponding parameter triples (*λ, μ, ρ*). Under the BD model with parameters (*λ, μ, ρ*), the density of an extant time tree is denoted *L*^(*λ,μ,ρ*)^(*τ*_1_, …, *τ*_*n*_). The precise form of *L*^(*λ,μ,ρ*)^ is not important for what follows, but can be found in e.g. Morlon et al. (2011, equation 1). Note that the topology of the time tree is uninformative in this model; the likelihood depends only on the merger times *τ*_*i*_

Abstracting away from the BD model, we now turn to the concept of identifiability. Let Θ be an arbitrary parameter space, and let *L*_*θ*_ denote a likelihood function parameterized by *θ* ∈ Θ. The *statistical model ℒ*_Θ_ = {*ℒ*_*θ*_: *θ* ∈ Θ} is the image of Θ under *L*_*θ*_; that is, the set of all possible likelihood functions that can be obtained from the parameter space Θ. (If Θ is a set of BD parameters, we use the notation

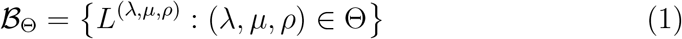

to emphasize that we are focusing specifically on the BD model.)

### Definition 1

(Identifiability). The statistical model ℒ_Θ_ = {*L*_*θ*_: *θ* ∈ Θ} is *identifiable* if *L*_*θ*_ is injective; that is, for all *θ*_1_, *θ*_2_ ∈ Θ, we have 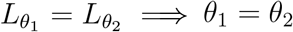

(In the BD model (1), the statement “(*λ*_1_, *μ*_1_, *ρ*_1_) = (*λ*_2_, *μ*_2_, *ρ*_2_)” is understood to mean that *ρ*_1_ = *ρ*_2_, and that the corresponding rate functions are equal almost everywhere.) If different parameters yield the same likelihood function, they cannot be distinguished using any amount of observable data. Identifiability is therefore the most minimal regularity condition one can place on a statistical model.

## 3 Results

In this section, we summarize LP’s results, prove that piecewise constant BD models are identifiable, and explore some additional corollaries and conjectures.

### 3.1 The result of Louca and Pennell

The key new quantity that underlies LP’s result is the so-called *pulled (birth) rate function*, which is defined as

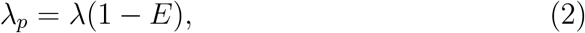

where *E*(*t*) is the probability that a lineage alive at time *t* has no descendants sampled at time 0. The antiderivative of *λ*_*p*_ is denoted

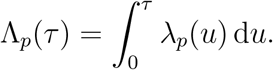

The function *E* satisfies the ordinary differential equation

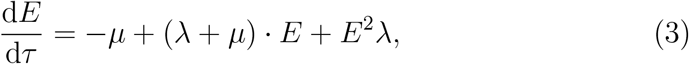

with initial condition *E*(0) = 1 − *ρ*. The solution to (3) is (Morlon et al., 2011)

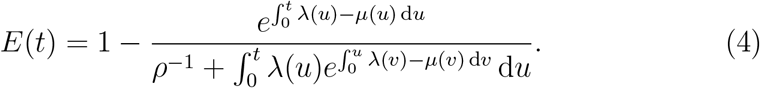

Note that *E*(*t*) is continuous, even if *λ* and *μ* are not.

The pulled rate function completely characterizes the likelihood of an extant timetree. Specifically, by equation (34) of LP,

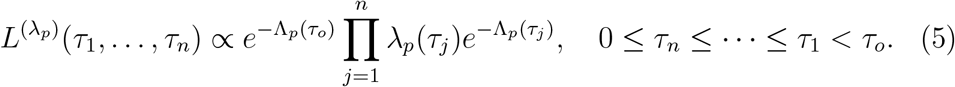

By implication, any two BD parameter triples (*λ*_1_, *μ*_1_, *ρ*_1_) and (*λ*_2_, *μ*_2_, *ρ*_2_) which generate the same *λ*_*p*_ via (2) are indistinguishable. LP‘s contribution is to show that this phenomenon emerges in a surprisingly general class of models. Restated in our notation, their main result is:

#### Theorem (LP)

*Given an extant timetree on n taxa with origin τ*_*o*_, *let* 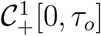 *denote the space of all functions which are strictly positive and continuously differentiable on* [0, *τ*_*o*_], *and let*

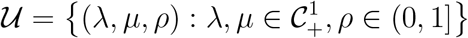

*be the set of all BD parameterizations derived from this space. Then the BD model* ℬ_U_ *is unidentifiable*.

Importantly, the theorem holds for any sample size *n*, and also if *ρ* is fixed. LP‘s proof is constructive and provides, for any given BD model, a set of infinitely many “congruent” models which all have the same likelihood. As LP argue in their discussion, this result has disturbing implications for the reliability of statistical estimates obtained from BD models, which have been widely reported in phylogenetics, phylodynamics, paleogenetics, and related fields.

### 3.2 Piecewise constant models are identifiable

In this section, we state and prove our main results.

#### Definition 2.

Let

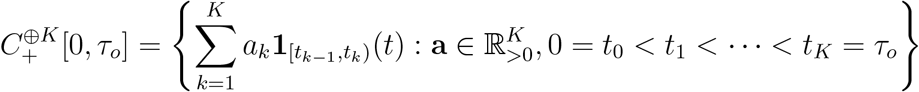

be the set of all positive piecewise constant functions with *K* pieces defined on [0, *τ*_*o*_].

Note from the definition that 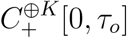 encompasses all possible piece-wise constant functions with *K* breakpoints; the breakpoints are not considered fixed or known in advance, and can vary from one function to another. Next, we define the class of BD parameterizations that forms the basis of our identifiability proof. In the definition and in what follows, we assume that the sampling fraction *ρ* ∈ (0, 1] is a fixed, known parameter. This is necessary because if *ρ* is allowed to vary, then as noted in the introduction, Stadler (2009) has shown that even the *constant rates* BD model is unidentifiable.

#### Definition 3.

Let

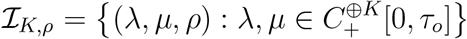

be the space of all piecewise-constant BD parameterizations with rate functions in 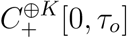 and fixed sampling fraction *ρ* ∈ (0, 1].

The following is our main result.

#### Theorem 4.

*If n >* 8*K, then the BD model* 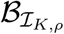 *is identifiable*.

The theorem is proved in Section 3.3. Unpacking the result, it asserts that *both* the positions of the breakpoints (the vector **t** in Definition 2) as well as the levels of each piece (the vector **a** in the definition) of both *λ*(*t*) and *μ*(*t*) are estimable given sufficient data, and that these breakpoints are not assumed to be shared between the two rate functions, or indeed between any two functions in the piecewise constant model space considered by the theorem. If, as is common in practice, we do assume that *λ*(*t*) and *μ*(*t*) are defined on the same set of breakpoints (while still allowing this set to vary between different parameterizations in ℐ_*K,ρ*_), then easy modifications to the proof show that *n >* 4*K* suffices for identifiability.

Note that the theorem only establishes a *sufficient* condition for identifiability. It does not imply that piecewise models are unidentifiable if *n* is below the stated bounds; in other words, we do not know if our bound is tight. In our view, the main message is that piecewise constant models are identifiable if 𝒪 (*K*) tips are sampled. Further implications of the theorem are discussed in Section 4.

Several extensions and conjectures follow naturally from Theorem 4. Since it is possible to uniformly approximate a regular function class over a compact set using step functions, identifiable BD models are in some sense dense in the space of all BD models. A prototypical result is:

#### Theorem 5.

*Let ρ* ∈ (0, 1] *be fixed, let*

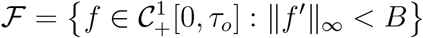

*be the set of positive, continuously differentiable functions with bounded first derivative over* [0, *τ*_*o*_], *and let* Θ_*ℱ,ρ*_ = {(*λ, μ, ρ*): *λ, μ* ∈ *ℱ} denote the resulting BD parameter space. Then*

1. 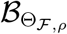 is unidentifiable; and
2. There exists a set of functions 𝒢 defined over [0, *τ*_*o*_] such that for any *ϵ >* 0,
  a. sup_*f* ∈ℱ_inf_*g*∈𝒢_ ∥*f* − *g*∥_∞_ *< ϵ*, and
  b. 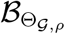 is identifiable if *n >* 8*Bτ*_*o*_*/ϵ*.

*Proof*. The first claim follows from LP, because their congruence classes include e.g. smooth perturbations of constant-rate BD models. For the second, if *f* ∈ ℱ then

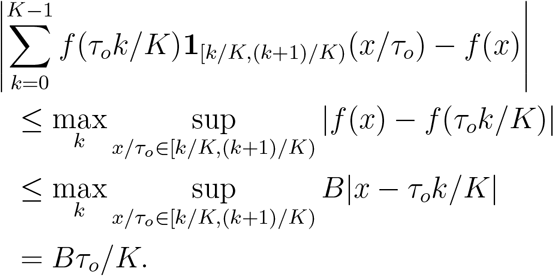

Letting *K* = *Bτ*_*o*_*/ϵ* yields the claim.□

An obvious caveat to this theorem is that the sample size needed to have (provable) identifiability grows rapidly as *ϵ* → 0.

Another possible extension relates to estimating birth-death models using polynomials. Since constant functions are polynomials of degree zero, it is natural to conjecture that identifiability holds for higher degrees as well.

#### Conjecture 6.

*Let* 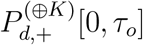 *be the set of non-negative, piecewise polynomials (splines) of order d with K* 1 *internal knots defined over* [0, *τ*_*o*_], *and* 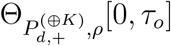 *be the corresponding BD parameter space, again for fixed ρ. Then the BD model* 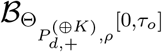 *is identifiable if n >* 8*K*(1 + *d*).

The conjecture would seem to imply that *n* grows with *d*, but this would be offset by having to use fewer pieces to obtain a good approximation. We are unable to prove this conjecture because substantial difficulties arise when trying to extend our proof technique to non-constant functions. Specifically, we do not know how to bound the sign change complexity of spline-based BD models (see Lemma 13) except when *d* = 0.

### 3.3 Proof of Theorem 4

Our proof derives from a general technique developed by Bhaskar and Song (2014) for establishing identifiability of rate functions in coalescent-type models. We follow their method closely, reproducing their results where necessary for completeness of exposition.

To build the necessary connections between the BD and coalescent models, we first notice that *L*^(*λ*^_*p*_) in (5) can be re-written as

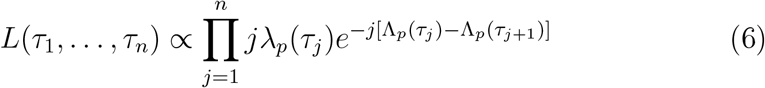

where we defined *τ*_*n*+1_ ≡ 0. This is a coalescent-type likelihood where the rate of merging (backwards in time) when there are *j* remaining lineages in the tree is *j*−1 instead of the usual 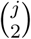. Hence, retrospectively, lineages are each killed at rate *λ*_*p*_(*τ*), except for one which must survive past time *τ*_*o*_.

Our strategy for establishing identifiability is to construct a vector of invariants which, for a sufficiently large sample size, uniquely identifies the pulled rate function *λ*_*p*_. To that end, given any pulled rate function *λ*_*p*_ and sample size *n*, we form an associated moment vector 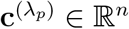, with entries

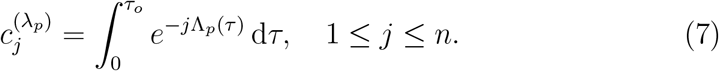

#### Proposition 7

*Suppose that* 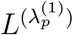 *and* 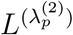 *are equal almost everywhere. Then* 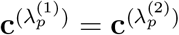.

*Proof*. In this proof we will refer to the likelihood function for multiple sample sizes, so we let 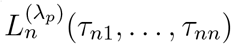 be the likelihood of an extant timetree with *n* + 1 sampled tips, where the merger times are *τ*_*o*_ ≥ *τ*_*n*1_ ≥ *τ*_*n*2_ ≥ · · · ≥ *τ*_*nn*_ ≥ 0. Expectation of a functional *f*: ℝ^*n*^ → ℝ with respect to 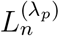 is denoted by 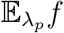; by definition, if 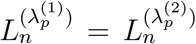 a.e., then 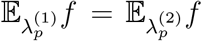 for all measurable *f*.

We use some results from Kamm et al. (2017) on moments of the truncated coalescent process, replacing each occurrence of the coalescent rate 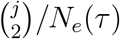 with its corresponding rate in the BD model, (*j* −1)*λ*_*p*_(*τ*). The expected value 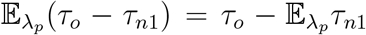 is written in their notation as 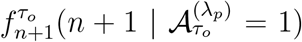, where 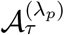 is the birth-death process analogue of the coalescent ancestral process, i.e. a pure death process on {*n* + 1, …, 1}, which begins at state *n* + 1 and transitions from state *j* + 1 to state *j* at rate *jλ*_*p*_(*τ*). By formulas (3) and (5) of Kamm et al. (2017), we have

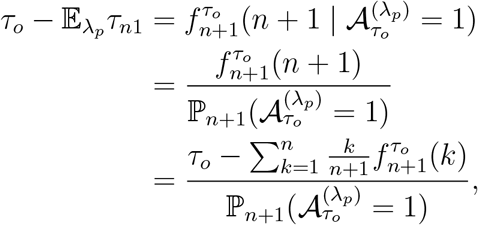

where 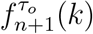 is defined below. The quantity 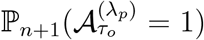 is the probability that the *unconditioned* birth-death process reaches a common ancestor before time *τ*_*o*_, meaning it is exactly the normalizing constant in equation (6). Rearranging the preceding display and defining 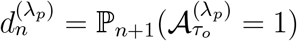, we obtain

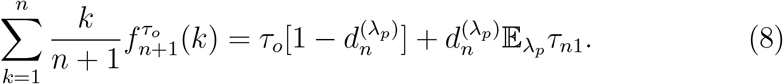

By Lemma 3.3 of Kamm et al., the summands in (8) are given by

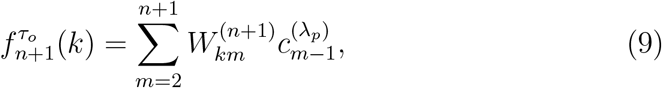

where the vector 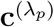 was defined in (7), and the matrix **W**^(*n*)^ ∈ ℚ ^(*n*−1)×(*n*−1)^ is defined by Polanski and Kimmel (2003, formula 10) to have entries

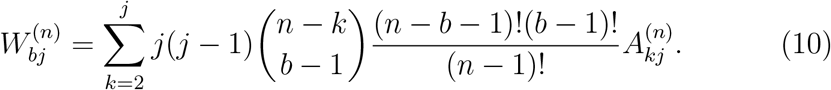

(Here, we follow the convention in the literature of indexing the second axis of **W**^(*n*)^ by 2, …, *n* instead of the usual 1, …, *n* − 1.) In the above display, the matrix **A**^(*n*)^ holds combinatorial coefficients which again have to be modified from their original definition (formula 6 of Polanski and Kimmel, 2003) to reflect the coalescence rate of the BD process:

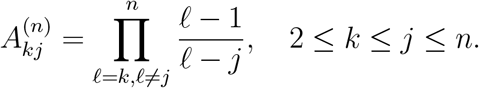

Now from (5), we have

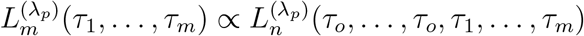

for any 1 ≤ *m* ≤ *n*. Thus, given any 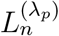, we may use the above procedure to calculate the moment vector

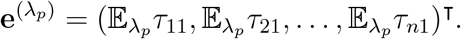

Define the lower-triangular matrix **B** = (*b*_*ij*_) ∈ ℝ ^*n*×*n*^ to have entries

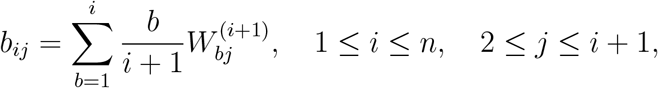

where the second axis of **B** is indexed in the same manner as **W**^(*n*)^. In Appendix A, we derive a closed-form expression for the entries of **B**, which shows in particular that the diagonal entries *b*_*i,i*+1_ = *i*(−1)^*i*+1^. Therefore, **B** is invertible, so that by (8),

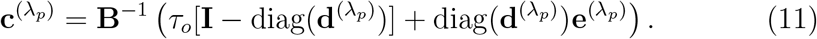

Finally, suppose that 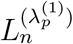 and 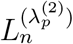 are two BD model likelihoods which are equal almost everywhere. Then there exists

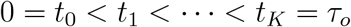

such that 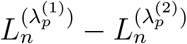 is continuous on open rectangles of the form

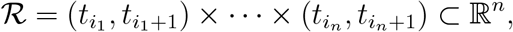

and equals zero almost everywhere on each such ℛ. Therefore, the pre-image

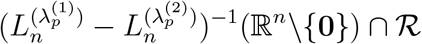

is an open set of zero measure; the only such set is ∅. Hence, 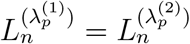 everywhere on ℛ. In particular, this implies that for all 1 ≤ *m* ≤ *n*, the BD likelihoods 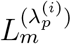 are equal almost everywhere on ℝ^*m*^. Therefore, the vectors 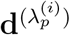 and 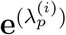, which are defined entirely in terms integrals of 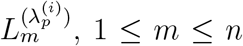, are equal for *i* = 1, 2. Equation (11) then implies that 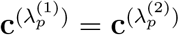.

Contrapositively, if 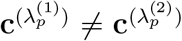, then 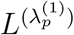 and 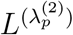 differ on a set of positive measure. The rest of the proof amounts to showing that if 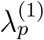 and 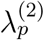 are generated by piecewise constant BD models, and *n* is sufficiently large, then they have different moment vectors.

The next theorem is restated for completeness.

#### Theorem

(Generalized rule of signs, Bhaskar and Song, 2014; Jameson, 2006). *Let f*: 𝒟 → ℝ *be a piecewise-continuous function defined on some domain 𝒟* ⊂ ℝ, *which is not identically zero and has a finite number σ*(*f*) *of sign changes. Then the function*

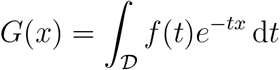

*has at most σ*(*f*) *zeros in* ℝ *(counted with multiplicity)*.

Informally, *f* is said to have a sign change any time it crosses zero, including by jump discontinuities. For a precise statement, refer to Definition 3 of Bhaskar and Song.

Given any pulled rate function *λ*_*p*_, we define its time-rescaled rate function

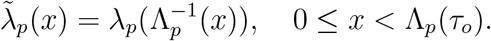

This transformation is invertible, since if

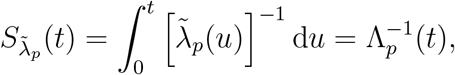

then 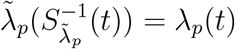. Hence,

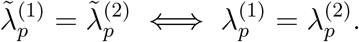

Then the entries of 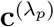 can be written as

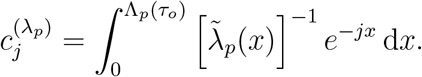

#### Proposition 8.

*Suppose that* 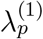 *and* 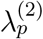 *are two pulled rate functions for which* 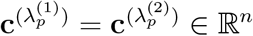. *Then either* 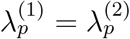, *or* 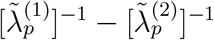 *has at least n* − 1 *sign changes over the shared domain of* 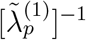 *and* 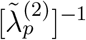.

*Proof*. Suppose that 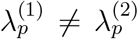. Then 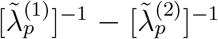 is not identically zero. Assume without loss of generality that 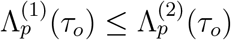 so the shared domain of 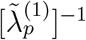 and 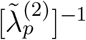 is 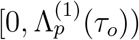. Consider the integral transform given by

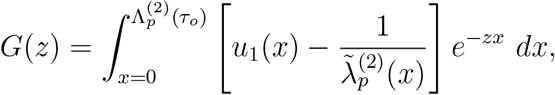

where

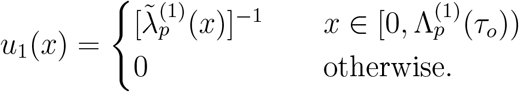

The supposition implies that *G*(*z*) has zeros at *z* = 1, …, *n*. By the generalized rule of signs, this implies that 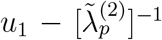 has at least *n* sign changes on 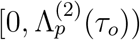. There is at most one sign change on the interval 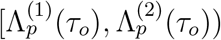, caused by a possible jump at 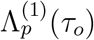. This implies that 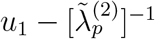 has at least *n* − 1 sign changes on 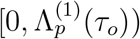.□

Based on the preceding result, we define the following complexity measure on BD model spaces. This is an adaptation of the Definition 4 in Bhaskar and Song to our setting. In the definition, the notation 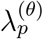 is used to denote the pulled rate function corresponding to a particular BD parameterization θ = (λ, μ, ρ).

#### Definition 9

(Pulled sign change complexity). Let Θ be a set of BD models.

The *pulled sign change complexity* 𝓁_p_(Θ) is defined as

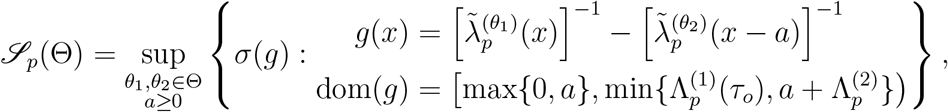

where σ(g) denotes the number of sign changes of g.

Using Proposition 8 and the preceding definition, we immediately have the following sample size criterion for the identifiability of BD models.

#### Theorem 10.

*Suppose that 𝓁_p_ (Θ) ≤ S, and that the mapping 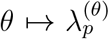 is injective over Θ. Then B_Θ_ is identifiable if n > S + 1.*

Theorem 10 is a general result that holds for any class of BD models. However, we can only apply it to spaces where the injectivity condition holds and where we know how to bound the pulled sign change complexity. Next, we establish these properties when Θ = ℐ_K,ρ_.

Recall that 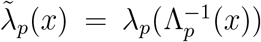 for x ∈ [0, Λ_p_(τ_o_)). By supplemental equation (9) of LP,

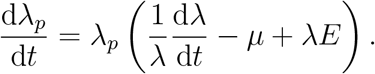

Hence,

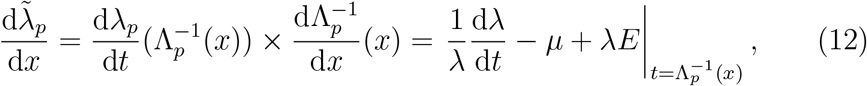

where in the second equality we used

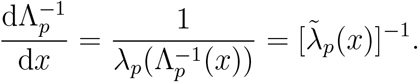

Now by (2),

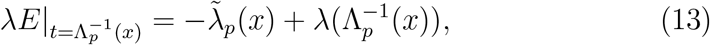

If *λ* and *μ* are constant, then d*λ*/d*t* = 0, and we obtain from (12) and (13) the first-order ordinary differential equation

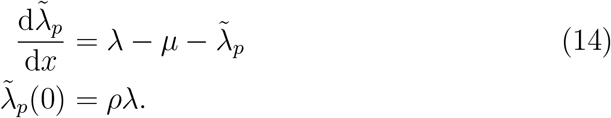

The solution to this differential equation is

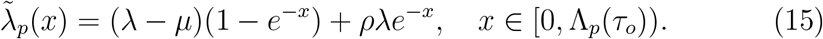

More generally, if *λ* and *μ* are constant over some interval [*t, t*^′^), then

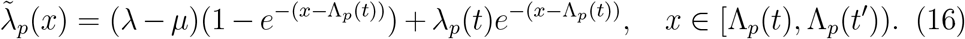

#### Proposition 11.

*Let θ_1_, θ_2_ be two different models in ℐ_K,ρ_ with pulled rate functions* 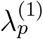 *and* 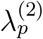. *Then* 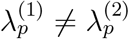.

Proof. Let (λ_1_, *μ*_1_) (λ_2_, *μ*_2_) be two different models in ℐ_K,ρ_. Then there is a non-empty interval [*t, t*^′^) ⊂ [0, τ_o_] such that

1. (λ_1_(*s*), *μ*_1_(*s*)) = (λ_2_(*s*), *μ*_2_(*s*)) for all 0 < s < t; and
2. λ_1_, *μ*_1_, λ_2_, *μ*_2_ are all constant over [*t, t*^′^) and (λ_1_(*s*), *μ*_1_(*s*)) ≠ (λ_2_(*s*), *μ*_2_(*s*)) for all t ≤ s < t^′^.

(Note that we could have *t* = 0, in which case condition 1 becomes vacuous.) To show 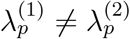, it is sufficient to show that 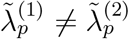. We assume that 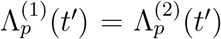, since if not the conclusion is immediate. By (16), for all 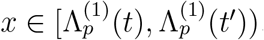, we have

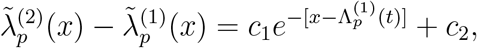

where

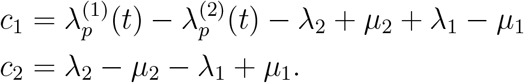

Suppose *c*_2_ = 0. Let *ε* = *E*^(1)^(*t*) = *E*^(2)^(*t*) ∈ (0, 1), where the equality follows from condition 1 and the facts that a) *E*(0) = ρ across all models, and b) *E*(*t*) is continuous (cf. equation 4). Then 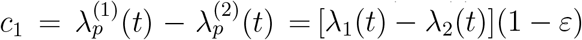. If *c*_1_ = 0 then this would contradict condition 2.□

*Remark*. The preceding result makes crucial use of the fact that all models in ℐ_*K*,ρ_ are constrained to have the same sampling fraction ρ. Without this assumption, the proposition would not even hold for *K* = 1 (Stadler, 2009).

Next, we bound 𝒫_*p*_(ℐ_*K*,ρ_). First, let

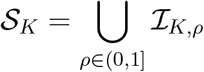

be the space of all *K*-piecewise constant BD models with unconstrained sampling proportions. As remarked above, this space is not identifiable, since in particular Proposition 11 does not hold for it. Nevertheless, it follows directly from Definition 9 that 𝒫_*p*_(ℐ_*K*,ρ_) ≤ 𝒫_*p*_(𝒮_*K*_), so bounding the pulled sign change complexity of _K_ is all that is required for our purposes.

We first show that 𝒫_*p*_(𝒮_K_) can be bounded in terms of the simpler quany 𝒫_*p*_(𝒮_1_).

#### Lemma 12.

*The pulled sign change complexity of 𝒮_K_ is bounded by the pulled sign change complexity of 𝒮*_1_ *as*

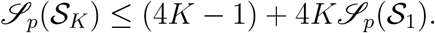

*Proof*. Let 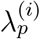 be the pulled rate function corresponding to (λ_*i*_, *μ*_*i*_, *ρ*_*i*_) for *i* = 1, 2. According to Definition 9, we need to bound all sign changes of

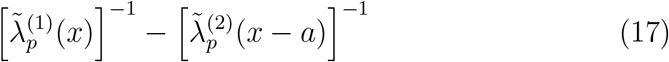

over the domain *x* ∈ [*m, M*), where

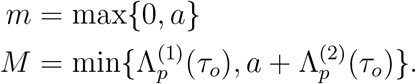

Enlarging the domain of (17) can only increase the number of sign changes, and the largest possible domain occurs when *a* = 0 and 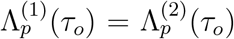 so we assume these conditions hold for the rest of the proof.

If 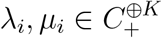, then we can place them onto a common set of 2*K* break-points

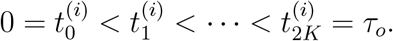

Let

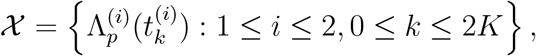

and sort the points in X to form a partition

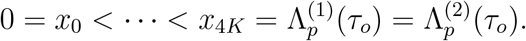

Allowing for possible jump discontinuities at *x*_1_, *x*_2_,…, *x*_4*K*_−_1_, the number of sign changes of (17) is at most 4K − 1 plus the number of sign changes on each interval (*x*_*j*_, *x*_*j*+1_).

For each i and j, there exists an integer 0 ≤ *k*(*i, j*) < 2K such that

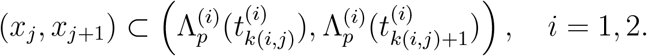

Therefore, there exists a BD parameterization θ_*ij*_ = (λ_*ij*_, *μ*_*ij*_, ρ_*ij*_) ∈ 𝒮_1_ such that

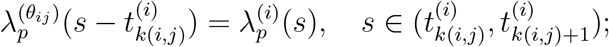

concretely, the initial sampling fraction is

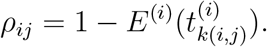

Then

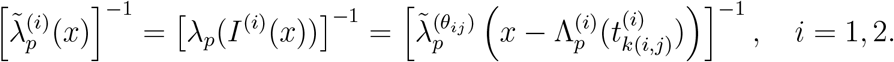

So within (*x*_*j*_, *x*_*j+1*_), the number of sign changes of 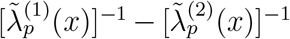 is at most the number of sign changes of

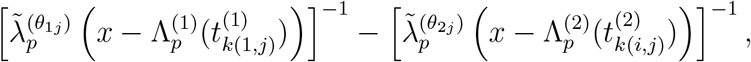

which is bounded above by 𝒫_*p*_(𝒮_1_). Hence, the number of sign changes is at most (4*K* − 1) + 4*K*𝒫_*p*_(𝒮_1_).

We conclude the proof by bounding 𝒫 (𝒮_1_).□

#### Lemma 13.

*Let (λ_1_, *μ*_1_, ρ_1_), (λ_2_, *μ*_2_, ρ_2_) ∈ 𝒮_1_, with corresponding pulled rate functions* 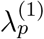 *and* 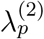, *and let*

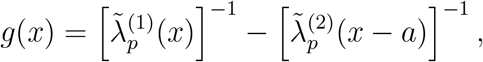

*where a ≥ 0 is arbitrary and the domain of *g* is as indicated in Definition 9. Then σ(*g*) ≤ 1.*

*Proof*. We have

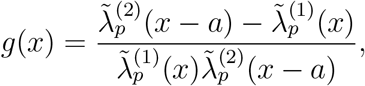

so the number of sign changes of *g* is at most the number of zeros of 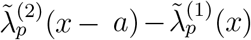. By (15), the function 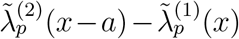 has the form c_1_e^−x^ +c_2_ for some *c*_1_, *c*_2_ that depend on λ_*i*_, *μ*_i_, ρ_*i*_, and *a*. Since this function is always monotone, 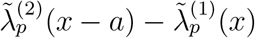 and hence *g* has at most one zero.□

By the preceding lemma, 𝒫_*p*_(𝒮_1_) ≤ 1, so that by Lemma 12,

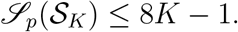

Finally, the claim follows from Theorem 10.

## 4 Discussion

In this paper, we proved that piecewise-constant BD models are identifiable from extant timetrees with a sufficient number of tips. We also showed that, under mild assumptions, unidentifiable BD models of the type considered by LP can be approximated to within arbitrary accuracy by identifiable BD models. Based on these results, we conjecture, but are unable to prove, that spline-based BD models are similarly identifiable.

In the short time since their publication, LP‘s findings have generated considerable discussion (e.g., Morlon et al., 2020; Helmstetter et al., 2021; Parry, 2021), with some authors concluding that they “will be dispiriting to evolutionary scientists” seeking to understand the factors affecting speciation and extinction (Pagel, 2020). Our results may serve to lift those spirits, while also illustrating potential subtleties that can arise when reasoning about a limiting concept like identifiability. For example, consider the BD models shown in Figure 2. The top row is reproduced from Figure 1 of LP and shows four color-coded BD models which all have the same pulled birth rate, and hence the same likelihood function. On the bottom, we approximated these functions over the domain [0, 16] using piecewise constant functions. By Theorem 4, these models can, in principle, be distinguished given a sufficiently large timetree. Is the underlying natural process that is modeled in Figure 2 inferable from data? The answer seemingly depends on whether the researcher believes that the piecewise functions shown on the bottom panel can faithfully represent this process. Empirically, we note that it would be nearly impossible to differentiate (using, say, a simple hypothesis test) between one of the 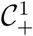 models in the figure and its corresponding 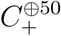 approximation on the basis of a realistically achievable amount of data.

**Figure 2:**
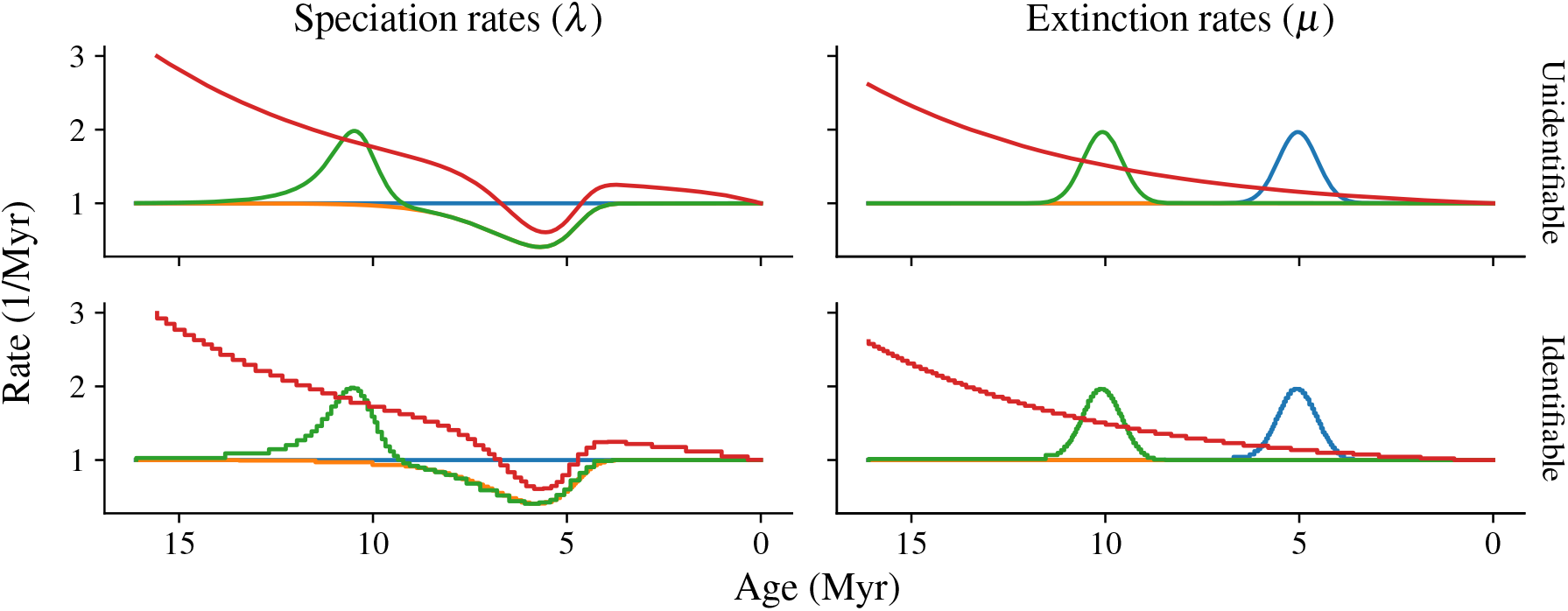
Unidentifiable vs. identifiable BD models. The top row contains four indistinguishable models exhibited in LP Figure 1. In the bottom row, we approximated these models using piecewise constant functions using *K* = 50 pieces. The models in the bottom row are identifiable given sufficiently many samples. All models are assumed to have the same *ρ*.

Still, practitioners should be careful not to overinterpret affirmative identifiability results as conclusive evidence that high quality estimates can be obtained on real problems. As we noted above, identifiability is a minimal regularity criterion that offers no guarantees about the ability to accurately infer model parameters using a finite amount of data. Even in identifiable models, it is often the case that potentially significant amounts of regularization and/or prior information have to be incorporated to obtain sensible results (Stadler et al., 2013; Kim et al., 2015; Bhaskar et al., 2015; Morlon et al., 2020; DeWitt et al., 2021). Having established identifiability, the next step would be to understand the finite-sample accuracy and rate of convergence of piecewise constant estimators in BD models. This is a challenging theoretical problem that will require new ideas and techniques. Fortunately, since several popular software packages (e.g., Bouckaert et al., 2019) already implement the piecewise constant BD model, there are already many simulation studies in the literature to help guide the way.

The reader may wonder whether our result is somehow a byproduct of the fact that we consider piecewise constant—hence discontinuous—rate functions, whereas in LP they are assumed to be continuously differentiable. In our opinion, this is not the main driver. Indeed, we believe that (cf. Conjecture 6) identifiable parameter spaces consisting of smooth functions also exist. Were it true, this conjecture would not contradict LP’s result, because the construction they use to generate their congruence classes (specifically, the operator *S*[*S*_o_, *f*] defined by supplementary equation 75) is not closed over simple function spaces like fixed-degree polynomials. In other words, even *f* is e.g. a spline, *S*[*S*_o_, *f*] is not. Thus, while there are infinitely large congruence classes of alternative BD parameterizations that are indistinguishable, the conjecture asserts that the intersection between these classes and a sufficiently simple function space consists of at most a single element. LP provide an heuristic argument supporting this conjecture in Section S.3 of their supplement.

Another criticism related to our use of piecewise constant models is that they require a potentially large number of samples to become (provably) identifiable. In some settings (e.g., phylodynamic analysis of a global pandemic) these bounds would be achievable, while in others they would plainly exceed the amount of available data. Using a more efficient representation like splines would lead to BD models with lower sampling requirements, but which currently lack identifiability guarantees.

In follow-up work, Louca et al. (2021) study a more general model where sampling is allowed to occur over time, and show that similar unidentifiability results hold in that setting as well. The coalescent-based methods we used in this paper, which condition on a number of lineages sampled at the present, do not readily extend to this setting, so our results leave open the question of whether piecewise-constant identifiability holds in random sampling models as well. In Section S.2.2 of their supplement, Louca et al. (2021) assert that restricting to piecewise constant model spaces cannot possibly resolve identifiability issues, however their argument is nonrigorous and based only on simulation evidence. Rather than posing a contradiction, we think their findings underscore the conclusions reached above: identifiability is fundamentally a mathematical property that cannot be checked on the basis of simulations alone, and it may have little bearing on one’s (in)ability to successfully carry out inference in real-world problems.

## Acknowledgments

We thank Andy Magee, Sebastian Prillo, and Ed Ionides for helpful comments on a draft of this manuscript. All errors are our own. This research was supported by NSF grants DMS-2052653 and DMS-1646108.

## A Computation of the matrix B

From the binomial identity

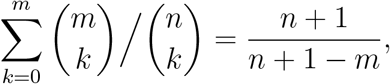

we obtain for k ≥ 2,

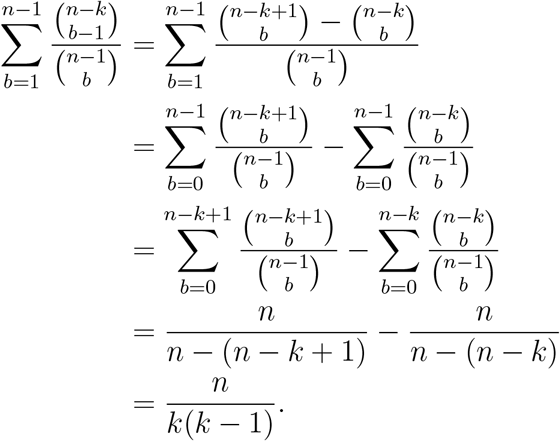

Hence,

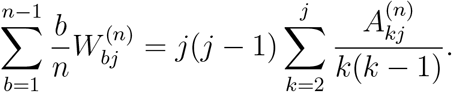

Furthermore,

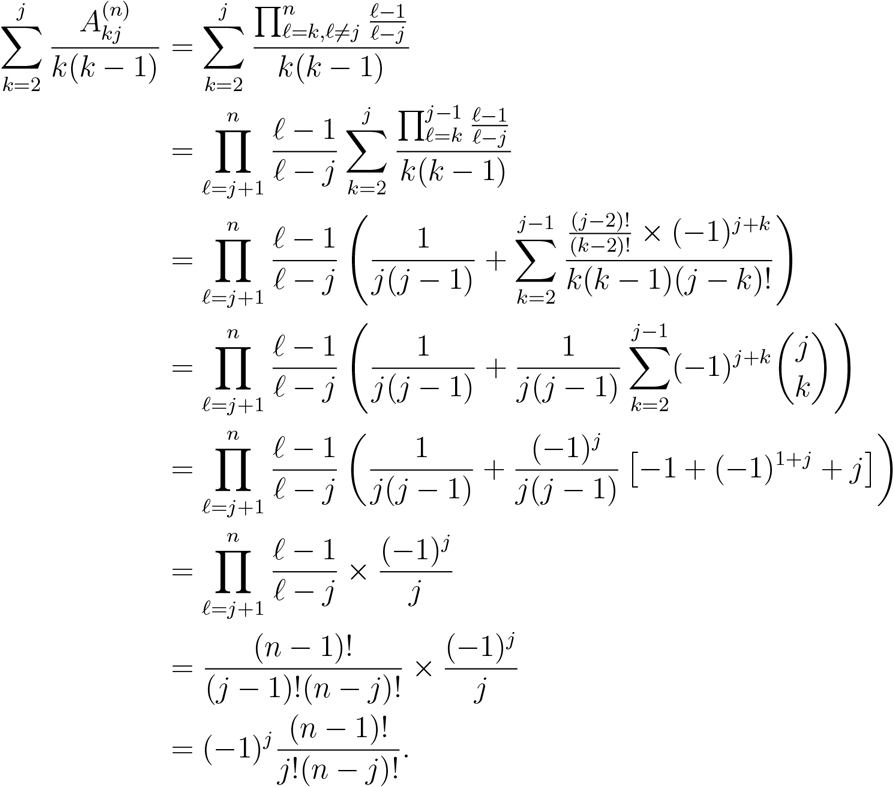

We conclude that

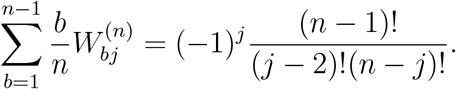

